# The bacterial endosymbiont *Wolbachia* increases reproductive investment and accelerates the life cycle of ant colonies

**DOI:** 10.1101/574921

**Authors:** Rohini Singh, Timothy A. Linksvayer

## Abstract

*Wolbachia* is a widespread group of maternally-transmitted endosymbiotic bacteria that often manipulates the reproductive strategy and life history of its solitary hosts to enhance its own transmission. *Wolbachia* also commonly infects eusocial insects such as ants, although the effects of infection on social organisms remain largely unknown. We tested the effects of infection on colony-level reproduction and life history traits in the invasive pharaoh ant, *Monomorium pharaonis*. First we compared the reproductive investment of infected and uninfected colonies with queens of three discrete ages, and we found that infected colonies had increased reproductive investment. Next, we compared the long-term growth and reproduction of infected and uninfected colonies across their life cycle, and we found that infected colonies had increased colony-level growth and early colony reproduction. These colony-level effects of *Wolbachia* infection seem to result because of a ‘live fast, die young’ life history strategy of infected queens. Such accelerated colony life cycle is likely beneficial for both the host and the symbiont and may have contributed to success of the highly invasive pharaoh ant.

## 1. Background

*Wolbachia,* a maternally-inherited group of endosymbiotic bacteria, infects an estimated 40% of arthropods [1,2]. It has a range of effects on host reproduction, including reproductive incompatibility between infected and uninfected mates, female-biased sex ratios in offspring of infected females [3,4], and increased fecundity of infected females [5,6]. These reproductive manipulations by *Wolbachia* can facilitate its spread in host populations, even when the manipulation is costly to the host itself [7–12]. Outside of reproduction, *Wolbachia* infection also has a spectrum of other phenotypic effects on its hosts, some of which can be beneficial in some context [3,4]. For example, *Wolbachia* infection alters the pheromonal profile of infected fruit flies. In the case of *Drosophila paulistorum,* this increases male mating success [13], whereas in the case of *D. simulans* this causes reproductive incompatibility [14]. These examples also show that *Wolbachia* can affect traits that influence social interactions in solitary species, suggesting that *Wolbachia* may similarly affect diverse individual- and group-level traits of highly social hosts such as ants.

An estimated 34% of ant species are naturally infected with *Wolbachia,* however its specific individual- and colony-level phenotypic effects remain unclear [15]. Given that *Wolbachia* often manipulates host reproduction to favor its own transmission [3–6,16–18], we studied the effects of infection on colony growth and reproduction dynamics in the pharaoh ant, *Monomorium pharaonis*. This ant species is one of the most successful and well-studied invasive ant [19]. It is polygynous (with multiple queens per colony), and colonies show natural variation in *Wolbachia* infection status [20,21].

To characterize the effects of *Wolbachia* infection on pharaoh ant colonies, we designed two separate assays. First we assessed the reproductive investment of infected and uninfected colonies that had queens of three discrete ages, namely 1, 3 and 6 months. In the second assay, we studied the life cycle dynamics of *Wolbachia* infected and uninfected colonies over a period of 7 months, spanning the entire life cycle of pharaoh ant colonies. Together, these assays compared growth, reproduction, and life history strategies of *Wolbachia*-infected and uninfected pharaoh ant colonies across the reproductive life span of queens.

## 2. Materials and methods

### (a) Source of experimental colonies and their infection status

Eight pharaoh ant lineages, originally collected from locations around the world were systematically intercrossed for nine generations in the lab to create genetically heterogeneous lab colonies [20–22]. Two out of these eight original lineages were infected by *Wolbachia* [20–22]. We identified the infection status of lab colonies using previously described PCR-based methods [23]. Specifically, we extracted genomic DNA from five individual workers per colony using Qiagen’s DNeasy Blood and Tissue kit (Cat. # 69506) by following the manufacturer’s protocol.

We mixed genetically heterogeneous lab colonies of the same infection status and evenly redistributed this mix to create 25 genetically homogeneous source colonies per infection group. We induced the production of new reproductives (queens and males) in these source colonies by removing the existing queens [24–28]. This synchronized the age of newly produced queens in these colonies. From this point on these colonies will be referred to as ‘queen age-matched source colonies’.We periodically examined and removed new reproductive larvae/pupae from these source colonies over the course of our experiments to ensure that queens in these colonies were the same age. All colonies were maintained and grown in environmental growth chambers at 27 ± 1°C, 50% RH and 12:12 LD cycle and were fed ad libitum synthetic agar diet (sugar:protein = 3:1) [29] and dried mealworms (*Tenebrio molitor*) twice a week.

### (b) Quantifying colony growth and reproduction dynamics

The pharaoh ant colony life cycle begins with intra-colony matings between newly produced males and queens (i.e. reproductives), followed by the production of only workers (growth phase), and ends with the spontaneous production of new reproductives (reproductive phase) when the existing queens senesce after approximately 7-8 months [30]. *Wolbachia* may manipulate different aspects of this colony life cycle to increase its transmission from one generation to the next. We designed two separate assays and compared the effect of *Wolbachia* infection on (a) colony-level reproductive investment in colonies with mated queens of known ages and, (b) colony life cycle dynamics (Fig. 1).

**Fig. 1.**
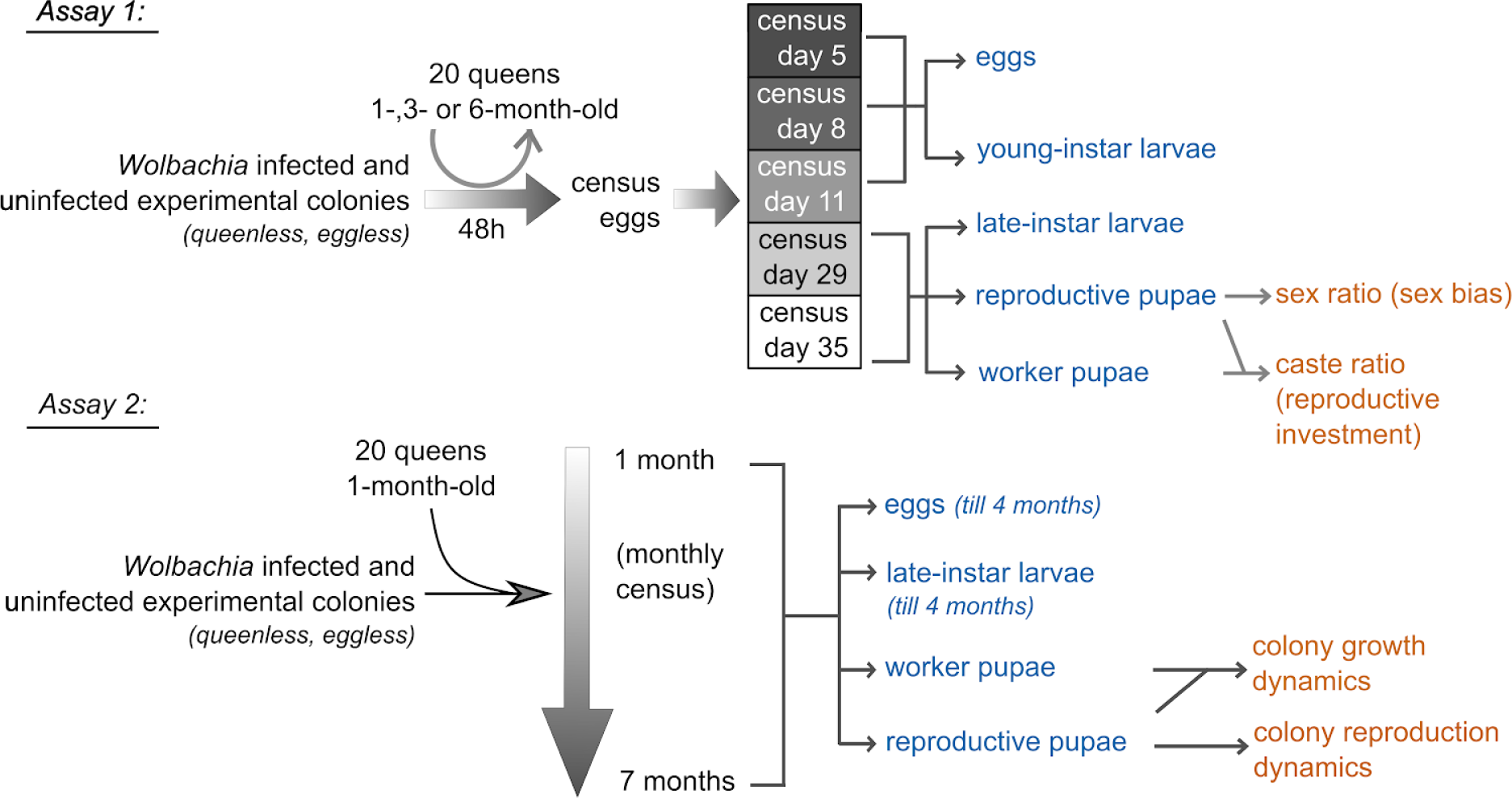
Schematic description of Assay 1 and Assay 2 for measuring the effects of *Wolbachia* infection status on productivity, reproduction, and life cycle of pharaoh ant colonies. We used Assay 1 (top) to assess colony-level reproductive investment at discrete queen ages and Assay 2 (bottom) to follow colony life cycle dynamics over time. We censused different ant development stages (in blue) at various times (arrows on the left of the development stages) to compute colony-level traits (orange) from various combinations of these census values (arrows on the right of the development stages).

#### Assay 1: Reproductive investment of colonies at discrete queen ages

We quantified reproductive investment of ten replicate infected and seven replicate uninfected colonies after inducing reproduction in these experimental colonies when queens in the colonies were 1-, 3- or 6-month-old. These ages span the reproductive lifetime of the queens.

We created experimental colonies for Assay 1 at the desired queen age by first mixing workers and brood (eggs, larvae, pre-pupae and pupae) from similarly infected lab-stock colonies and queen age-matched source colonies. We then redistributed ∼500 workers and ∼500 brood from this mix to create ten infected and seven uninfected experimental colonies of similar size. These experimental colonies were reared without queens for ten days to make the colonies eggless (Fig. 1). Once eggless, 20 age-matched mated queens from source colonies were added to these experimental colonies for 48h to produce developmentally synchronized batch of eggs (Fig. 1). After 48h of adding queens to these experimental colonies, we counted the total number of eggs present in these colonies and recorded the initial composition of the colony. After census, we removed the queens and returned them to their respective queen age-matched source colonies (Fig. 1). Over the subsequent five weeks, eggs transitioned to worker, male, and queen pupae, and we censused these pupae after 29 and 35 days of of adding the queens (Fig. 1). For each type of pupae, we calculated productivity as the sum of pupae counts on day 29 and day 35. Using these counts, we calculated relative investment in reproduction or colony caste ratio as the proportion of queens produced per total number of new females, and colony sex ratio as the proportion of queens produced per total number of new reproductives [20].

#### Assay 2: Colony growth, reproduction, and life cycle dynamics

We tracked colony growth and reproduction of 14 infected and 12 uninfected experimental colonies for seven months in order to quantify effects of *Wolbachia* on colony productivity, of both new workers and new reproductives, as well as to characterize the effects of *Wolbachia* on the colony life cycle.

For Assay 2, we created queenless and eggless experimental colonies in the same manner as described for Assay 1. Once eggless, we added 20 one-month-old mated queens from the queen age-matched source colonies to each experimental colony of the same infection status (Fig. 1). We censused the colonies after 48h to quantify the initial colony composition and did not manipulate the colonies any further. The queens aged naturally in these colonies and we surveyed colony composition across the whole colony life cycle. Specifically, we quantified colony growth and reproduction on a monthly basis for the first four months, by counting different developmental stages, from eggs to pupae, and reproductive adults (Fig, 1). After four months, the colonies were sizeable and it was difficult to get accurate counts of younger developmental stages. Hence, after four months we restricted the counts to reproductive (i.e. new males and queens) adults and pupae, and worker pupae (Fig, 1). At each time point, we calculated net productivity as the total number of pupae (workers, queens and males) present at the time of census (Fig, 1).

We also collected 15 white colored worker pupae, with pigmented eyes, after 2, 3, 4 and 6 months each of starting the assay. We dried these pupae at 55°C for 20h before storing them at −20°C till the time of weighing them on Sartorius microbalance (MSU3.6P-000-DM) in milligrams up to three decimal points.

### (c) Statistical analysis

We used R version 3.5.2 [31], with lme4 [32], pscl [33], MASS [34] and car packages [35] for data analysis, and ggplot2 [36] for plotting graphs. We constructed generalized linear mixed effect models (GLMM; [37]) to assess the effects of predictor variables (*Wolbachia* infection and queen age or time) on response variables (fitness traits such as total number of queens, sex ratio, and caste ratio), with queen age-matched source as a random factor. To assess the effect of *Wolbachia*-by-queen age (Assay 1) or *Wolbachia*-by-time (Assay 2) interaction on fitness traits, we used generalized linear models (GLMs; [37]) with *Wolbachia* infection, queen age/time and *Wolbachia*-by-queen age/time interaction as fixed factors. We also used GLMs to assess the effect of *Wolbachia* on fitness traits at specific queen ages/time points. For count data, we constructed GLMMs with Poisson and GLMs with negative binomial or quasi-Poisson error distributions. For caste and sex ratio, we constructed GLMMs assuming binomial and GLMs assuming quasi-binomial error distributions. For Assay 2, we split the analysis in two parts. For the first part we used data from one to four months, including late-instar larvae as a fixed factor, and for the second part we used data from five to seven months which did not include late-instar larvae counts. For colony reproduction in Assay 2, we used counts of queen and males from four to seven months for analysis. For dry mass, we used linear mixed effect models (LMM; [38]) with *Wolbachia*-by-time interaction term as predictor variable, log-transformed dry mass as the response variable, and experimental colonies as random factor. For age-specific effects of *Wolbachia* infection, we constructed LMM as described above, with *Wolbachia* as predictor variable. Datasets for Assay 1 and Assay 2, and R scripts have been included as supplementary information (Appendix S1-S5).

## 3. Results

### (a) Wolbachia increased queen production and reproductive investment of colonies with reproductively mature queens

Overall, *Wolbachia*-infected colonies had increased reproductive investment since they produced more queen pupae (GLMM; LRT = 8.75, *p* = 0.003; Fig. 2a) and had higher queen-biased caste ratio (GLMM; LRT = 5.88, *p* = 0.015; Fig. 2b), specifically when infected colonies had 3-month-old queens (total number of queen pupae, GLM: *F* = 5.63, *p* = 0.031 and caste ratio, GLM: *F* = 9.01, *p* = 0.009; Fig. 2). However, infected and uninfected colonies produced a similar number of males (GLMM; LRT = 0.03, *p* = 0.84; Fig. S1b) and had a similar colony-level sex ratio (GLMM; LRT = 2.71, *p* = 0.09; Fig. S1c). In addition to *Wolbachia* infection, queen age also affected colony-level traits. The total number of eggs present in the experimental colonies after 48h increased with queen age (GLMM; *F* = 1421.15, *p* < 0.001; Fig. S2a). The total number of queens produced from these eggs was also dependent on queen age (GLMM: LRT = 419, *p* < 0.001), with increased production of queen pupae in experimental colonies with 3-month-old queens (GLM: z < 18, *p* < 0.001; Fig. S2b). Furthermore all colonies with older queens produced more males (GLMM: LRT = 197.63, *p* < 0.001; Fig. S2c), had male-biased sex ratios (GLMM: LRT = 122.2, *p* < 0.001; Fig. S2e), and had worker-biased caste ratios (GLMM: LRT = 571.11, *p* < 0.001; Fig. S2f)

**Fig. 2:**
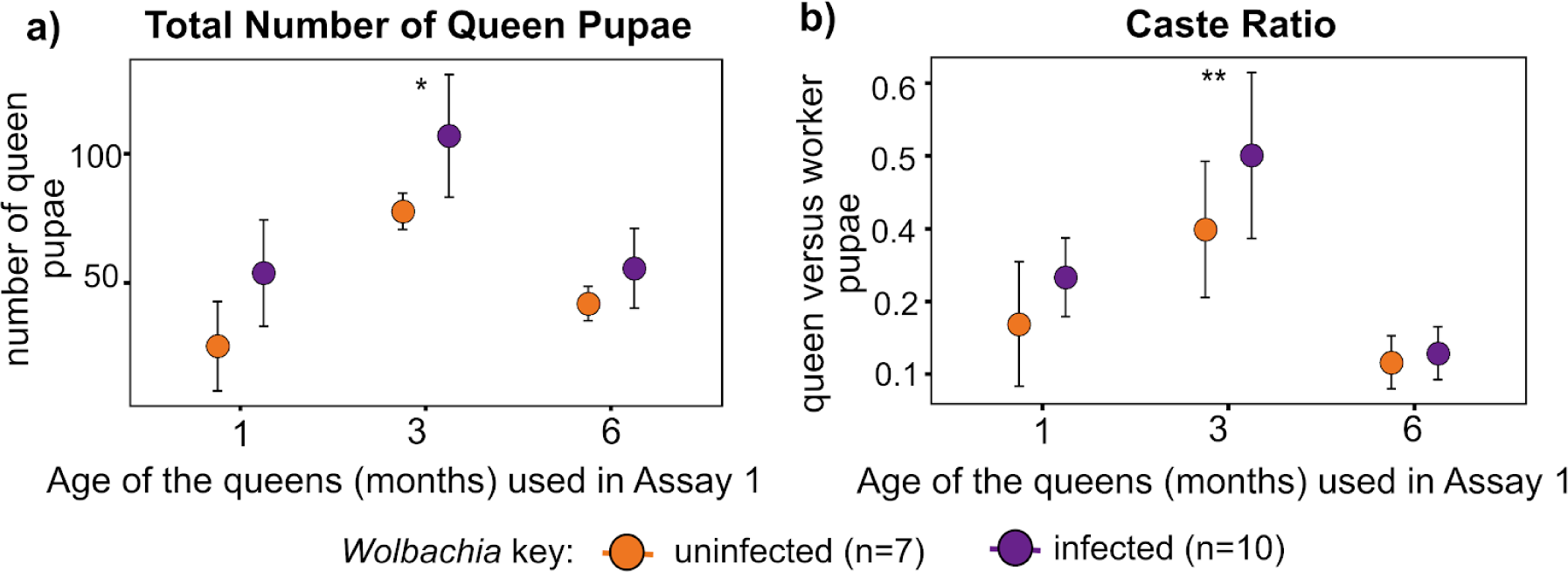
*Wolbachia* increases reproductive investment of pharaoh ant colonies, depending on queen age. (a) Infected colonies produced more queen pupae when queens used for the assay were 3-month-old. (c) Infected colonies have increased queen-biased caste ratio when queens used for the assay were 3-month-old. Filled circles represent the mean trait value and error bars represent the 95% confidence interval of the mean. *Wolbachia*-driven differences are represented as *p* < 0.05* and <0.01**, and were estimated by age-specific GLMs. The number (n) of replicate colonies in the assay are at the bottom of the figure panel.

In summary, our results for Assay 1 show that *Wolbachia* increased colony-level reproductive investment of infected colonies, specifically of colonies with 3-month-old queens. Furthermore, queen age, independent of *Wolbachia,* was the primary predictor of colony-level productivity differences.

### (b) Infected colonies have increased colony-level growth, early colony reproduction and faster colony life cycle

Similar to our results from Assay 1, colony-level productivity traits of *Wolbachia*-infected colonies were different from uninfected colonies only at certain time points. Infected and uninfected colonies produced a similar number of eggs (GLMM: LRT = 0.4, *p* = 0.51), although the number of eggs consistently increased in all the colonies over time (GLMM: LRT = 1232.2, *p* < 0.001; Fig. S3a). Infected colonies had more late-instar larvae, particularly after two months of starting the assay (GLM: *F* = 4.85, *p* = 0.039; Fig. S3b). Furthermore, the total number of worker pupae produced by infected colonies between five and seven months was dependent on time point (GLM: LRT = 4.22, *p* = 0.018), with infected colonies producing more worker pupae after two months (GLM: *F* = 7.6, *p* = 0.012; Fig. 3a), six months (GLM: *F* = 6.4, *p* = 0.019; Fig. 3a) and seven months (GLM: *F* = 6.38, *p* = 0.019; Fig. 3a) of starting the assay. Similarly, the dry mass of infected worker pupae was dependent on time (LMM: *X*^*2*^ = 153.13 *p* < 0.001; Fig. S4), and infected worker pupae were heavier after two months of starting the assay (LMM: *X*^*2*^ =8.69, *p* = 0.003; Fig. S4).

**Fig. 3:**
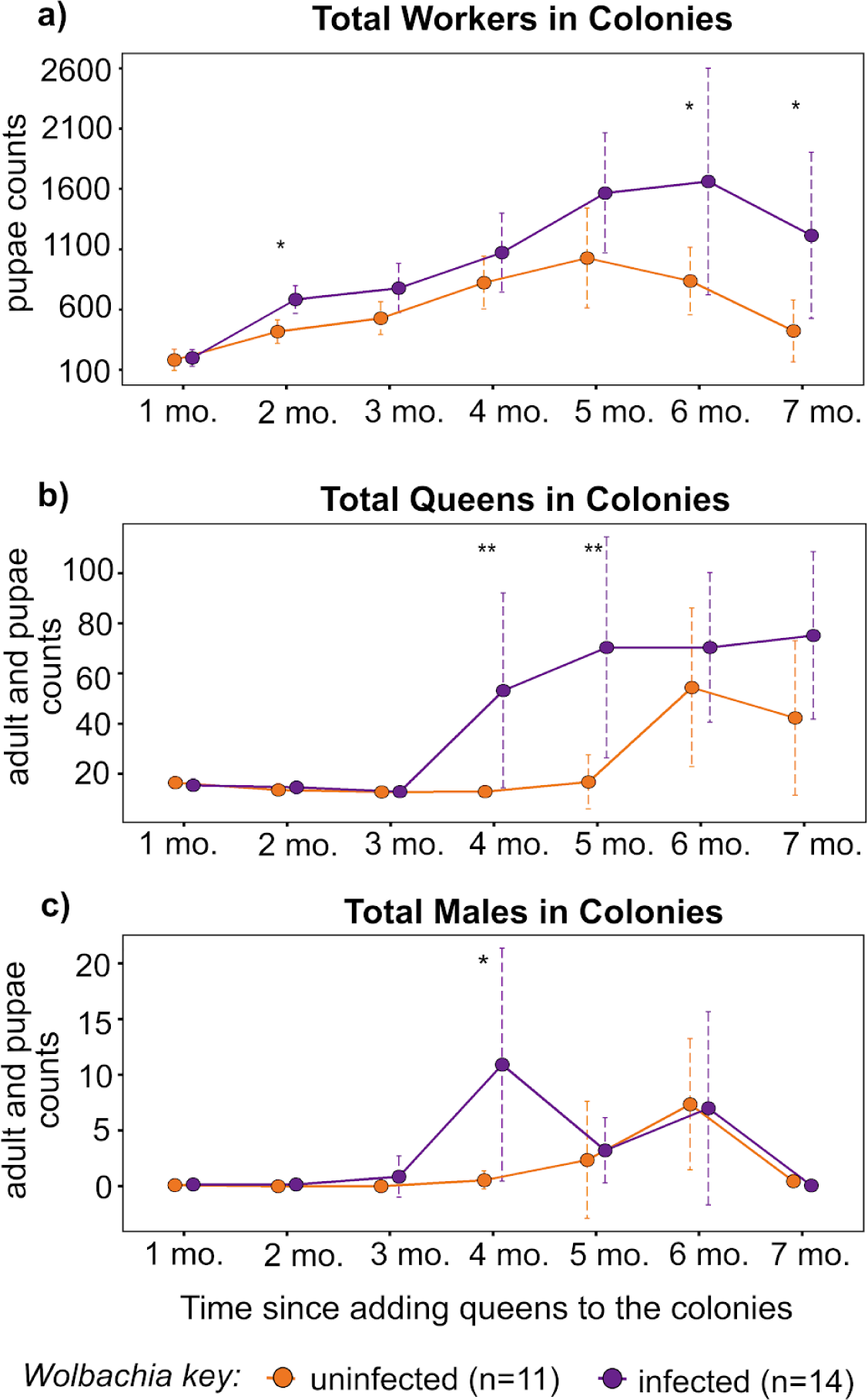
Infected colonies had increased growth and early onset of reproduction. (a) Infected colonies produced more pupae at two months after starting the assay. (b) Infected colonies had an early spontaneous production of new queens. (c) Infected colonies had an early spontaneous production of new males. Filled circles represent the mean trait value and error bar represents the 95% confidence interval of the mean. *Wolbachia*-driven differences are represented as *p* < 0.05* and <0.01**, and were estimated by age-specific GLM. The number (n) of replicate colonies in the assay are at the bottom of the figure panel.

Infected colonies had more queens after four months (GLM: *F* = 8.5, *p* = 0.007) and five months (GLM: *F* = 12.44, *p* = 0.002; Fig. 3b) of starting the assay, and had more males after four months of starting the assay (GLM: LRT = 7.81 *p* = 0.02; Fig. 3c). Since infected colonies reproduced earlier, this suggests that *Wolbachia* infection accelerated colony life cycle dynamics.

## 4. Discussion

The effects of *Wolbachia* on reproduction and physiology of solitary species are well-studied [4]. However, despite its wide occurrence in ants, the effects of infection on social life are currently unknown. We provide the first evidence for effects of *Wolbachia* on the life history strategy of ant queens, and reproductive investment and life cycle of ant colonies.

We show that *Wolbachia*-infected pharaoh ant colonies have a reproductive (Fig. 2) and growth (Fig. 3a) advantage that is dependent on the age of the queens. Furthermore, infected colonies shift from exclusively producing workers to producing new reproductives (i.e. new queens and males) earlier than uninfected colonies. This suggests that infected queens experience early reproductive senescence, since the presence of reproductively fecund queens in pharaoh ant colonies suppresses the production of new queens and males [24,25,28,30]. This accelerated reproductive senescence could potentially arise due to a trade-off between fecundity and somatic maintenance in the infected queens [39,40]. While such a trade-off fundamentally occurs at the level of individual queens, it affects collective decisions for colony reproduction. Our results point to an alternate ‘live fast, die young’ life history strategy of infected queens, which acts to accelerate the colony life cycle. Furthermore, our results also underscore the importance of queen age on colony life cycle dynamics.

An accelerated ant colony life cycle will act to increase the frequency of colony reproduction (i.e. decrease the generation time) of infected relative to uninfected colonies, which will especially be favored in expanding populations. Invasive species such as pharaoh ants likely find themselves in conditions where such rapid population expansion is favored, e.g., following invasion into a new habitat. New pharaoh ant colonies are established when some of the existing queens and workers “bud” off from the parent colony and occupy new nest sites [30,41]. Our results suggest that *Wolbachia* infection may increase the frequency of such colony-founding events and hence, increase the invasiveness of infected pharaoh ant colonies. Given the growth advantage of infected colonies, *Wolbachia* infection may be expected to sweep through pharaoh ant populations, as has been shown previously for solitary host species [11,42].

The probability of infection sweeping through ant populations and increasing the invasiveness of infected populations, will however depend on multiple factors. Ant colony growth and reproduction is socially regulated [27,43–47], and hence the spread of *Wolbachia* can be limited by intra-colony as well as inter-colony interactions. Rapidly expanding invasive pharaoh ant colonies will likely come in contact with both infected and uninfected colonies. Pharaoh ant colonies show little inter-colony aggression, and colonies in the laboratory readily merge despite being highly genetically differentiated, but it is uncertain how frequently and readily colonies merge in nature [22]. Future studies investigating the within-colony dynamics of *Wolbachia* infection will further elucidate how *Wolbachia* is expected to spread across colonies and populations.

Interestingly, we did not observe reduced male production or queen-biased sex ratios in infected colonies, as we did in a previous study using artificial selection on pharaoh ant caste ratio for three generations [20]. However, we observed increased queen production and queen-biased caste ratios, which similarly results in relatively increased investment in female reproductives. Thus, both studies point to reproductive manipulation by *Wolbachia* that is expected to increase its own transmission to the next generation. Such colony-level fitness effects of *Wolbachia* infection, while similar to the effects observed in solitary species, are expected to partly result from mechanisms fairly unique to social organisms. For example, infected pharaoh ant colonies produced more pupae while producing a similar number of eggs compared to uninfected colonies (Fig. 3a, Fig.S3a respectively). This suggests that infected colonies have a higher egg-to-pupa survival. This could be attributed to individual-level differences in the quality of the eggs laid by the queens or the collective differences in foraging and nursing behaviors of infected workers, or both. *Wolbachia* is a nutritional mutualist in other species [48–50]. It is also possible that *Wolbachia* could be supplementing ant queens in a similar manner, which may increase the nutritional quality and survival of their eggs. Further studies are necessary to tease apart the specific individual- and colony-level mechanisms underlying the effects of *Wolbachia* that we have observed.

In summary, we show novel colony-level fitness and life history effects of a widespread insect endosymbiont in a social organism. *Wolbachia* infection profoundly altered the life history strategy of queens and accelerated the colony life cycle of pharaoh ants. This effect of *Wolbachia* on its ant host may have evolved as a means to increase its own transmission. At the same time, under the environmental conditions of our study, and likely under conditions commonly experienced by invasive pharaoh ants, these effects are beneficial to the host as well.

## Author contributions

All authors conceived the study. RS collected and analysed the data, and drafted the manuscript. TAL critically revised the manuscript.

## Acknowledgements

We thank Isaias Jacinto (University of Pennsylvania), Naima Okami (Brown University) and Matthew Dougherty (University of Pennsylvania) for assisting with assay 1 and assay 2 at various time points. We are grateful to Chao Tong, Michael Warner and Justin Walsh for their feedbacks at various steps of the current work and for their comments on multiple drafts of this manuscript. We are extremely grateful to Jacob Russell (Drexel University) for critical inputs at different stages of this work.

