## Supplementary figures S1-S5 for "The bacterial endosymbiont *Wolbachia* increases reproductive investment and accelerates the life cycle of ant colonies"

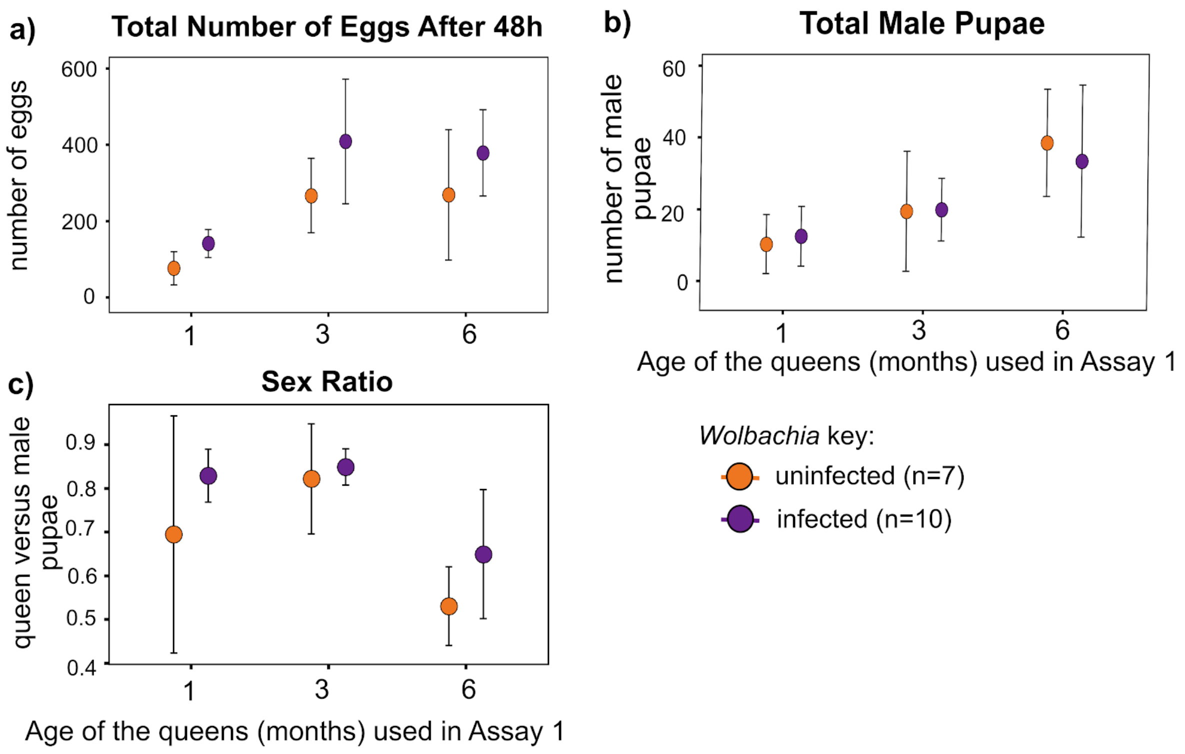


**Fig. S1: No differences between infected and uninfected colonies for some colony-level fitness traits.** (a) Total number of eggs in experimental colonies after adding 20 age-matched queens. (b) Infected and uninfected colonies produced similar number of males. (c) No differences between sex ratio of infected and uninfected colonies. X- axis represents the discrete queen ages used in Assay 1, Y-axis represents the trait value, filled circles represent the mean trait value and error bar represents the 95% confidence interval. Sample sizes per group (n) are provided with color key for *Wolbachia* infection status of colonies.

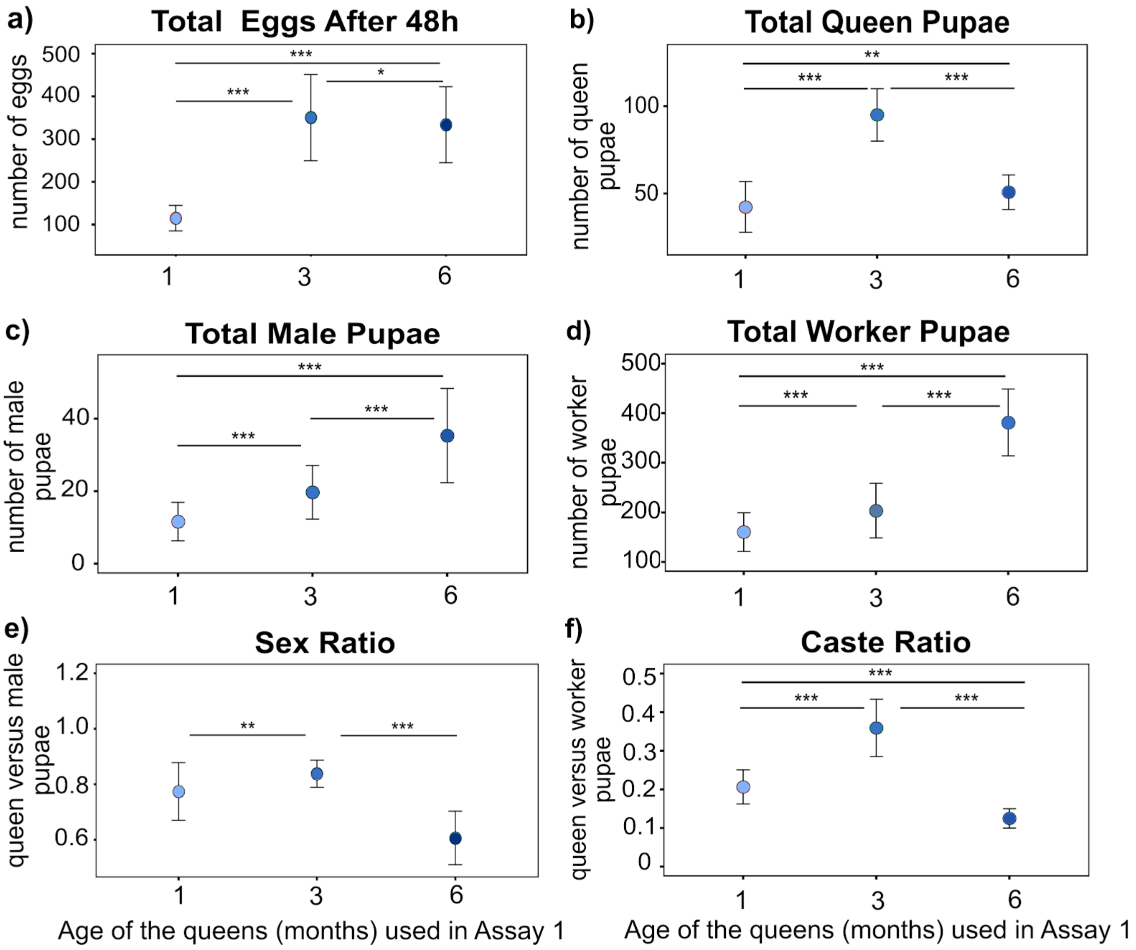


**Fig. S2: Colony-level fitness traits vary across queen age.** (a) One month-old queens laid the least number of eggs within 48h. (b) Colonies with three months-old queens produced the highest number of queen pupae. (c) Male production increased as the queens became older. (d) Worker production increased as the queens became older. (e) Male biased sex ratio in older queens. (f) Colonies with three months-old queens had the higher queen-biased caste ratio. X- axis represents the discrete queen ages used in Assay 1, Y-axis represents the trait value, filled circles represent the mean trait value and error bar represents the 95% confidence interval. Statistical differences, as estimated by TukeyHSD of GLMM for effect of queen ages, are represented by *p < 0.05, **p<0.01 and ***p<0.001. 16 colonies were analyzed per time point.

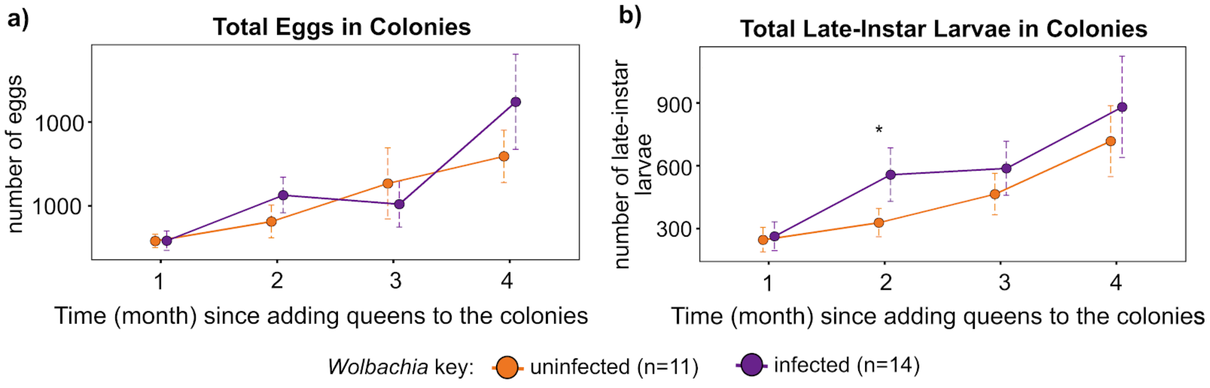


**Fig. S3: Growth dynamics of the early developmental stages in colonies.** (a) Infected and uninfected colonies produced similar number of eggs. (b) Infected colonies had higher number of late-instar larvae after 2 months of adding queens to experimental colonies. X-axis represents the time, in months since Assay 2 was started, Y-axis represents the trait value, filled circles represent the mean trait value and error bar represents the 95% confidence interval. *Wolbachia*-driven difference is represented as *$p < 0.05$, and was estimated by age-specific GLM. *Wolbachia* color key, along with the number of colonies in the assay (n), are at the bottom of the figure panel.

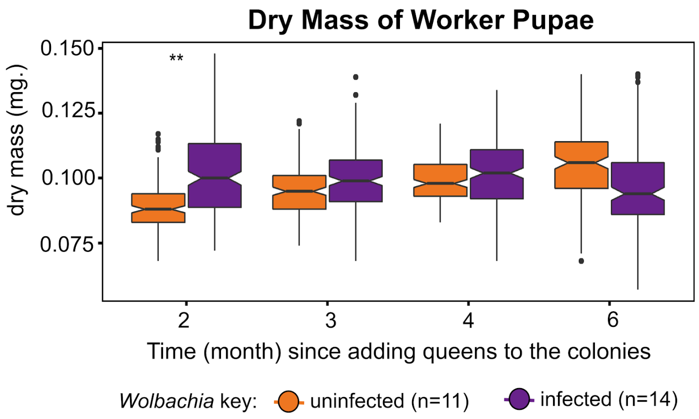


**Fig. S4: Dry mass of worker pupae varies across queen age.** Infected worker pupae were heavier after 2 months of starting Assay 2. X-axis represents the time, in months since Assay 2 was started, Y-axis represents the trait value, *Wolbachia*-driven differences are represented as **p<0.01, which was estimated by ANOVA of age-specific LME. *Wolbachia* color key, along with the number of colonies (n) in the assay, are at the bottom of the figure panel.

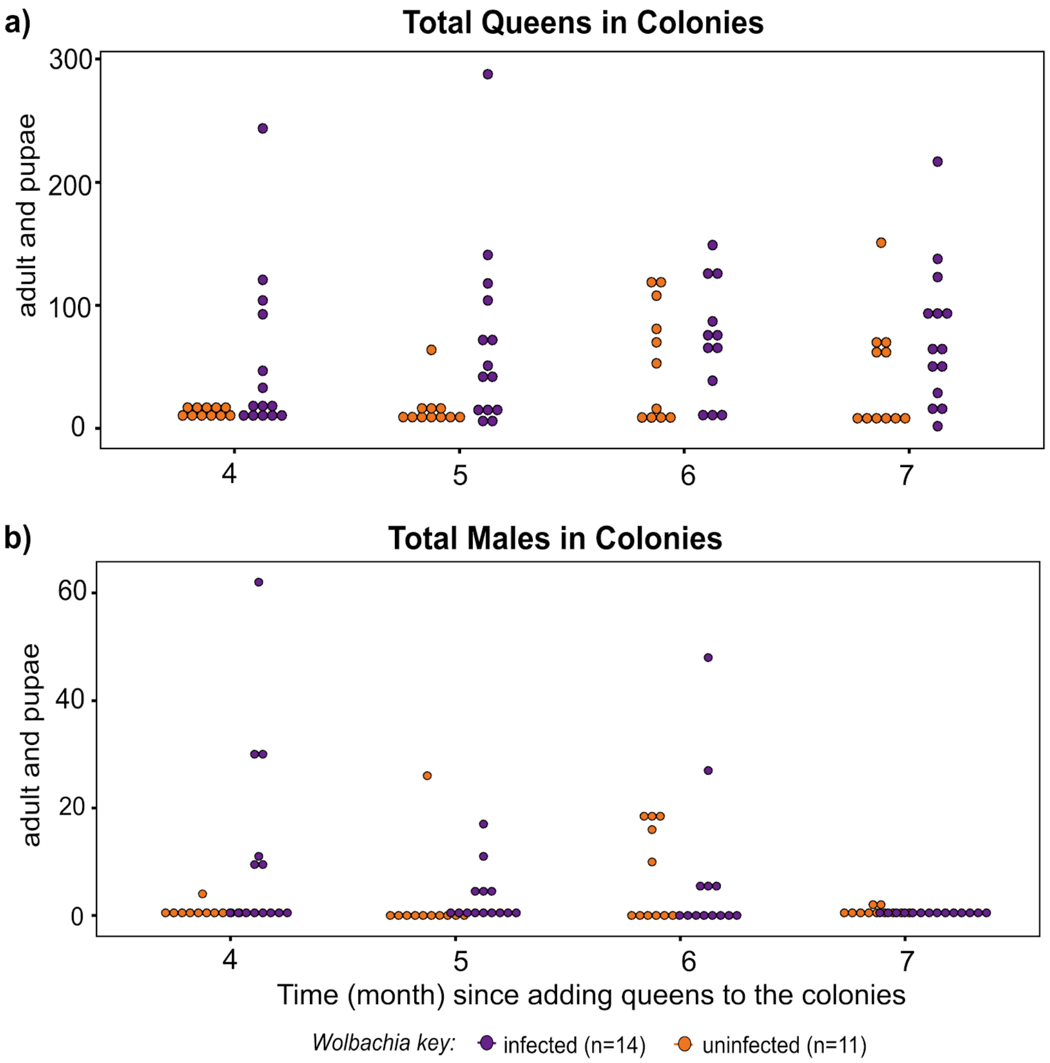


**Fig. S5: Early onset of reproduction in infected colonies.** X-axis represents the time, in months, since the assay was started by adding 20 one month-old queens in experimental colonies and Y-axis represents the trait value, filled circles represent the raw trait value. *Wolbachia* color key, along with the number of colonies in the assay (n), are at the bottom of the figure panel.
