## Appendix S5 for "The bacterial endosymbiont *Wolbachia* increases reproductive investment and accelerates the life cycle of ant colonies"

#-------------------- GOAL: COLONY LIFE CYCLE DYNAMICS OVER TIME --------------
# loading required packages ----
library("lme4") #LM, LMM, GLM, GLMM
library("dplyr")
library("car") #anova for LM and LMM
library ("MASS") #negative binomial GLM
library ("pscl") #hurdle and zero-inflation models
library("multcomp")
library("Rmisc") #stats (not correlation)
library("ggplot2") #for all graphs


# ----------------- Loading files and processing it before analysis ------
# Loading files and creating variable -------
setwd("G:/My Drive/Linksvayer_wolprojects/manuscripts/1_fitness effects of Wolbachia/supp_inf")
list.files() #visualize files in the folder
dm <- read.csv("raw_data_assay2_mass.csv")
lt <- read.csv("raw_data_assay2_census.csv")

#setting something as a factor
lt$exp_code = as.factor(lt$exp_code)
lt$nests = as.factor (lt$nests)

# creating new variables for measuring colony productivity and reproductive output
lt$tot_male = lt$sex_pup_male + lt$males #total males in colonies at a time point
lt$tot_qn = lt$sex_pup_fem + lt$queens #total queens in colonies at a time point
lt$prod = lt$worker_pup + lt$sex_pup_male + lt$sex_pup_fem #using only pupae counts for net productivity

#saving the above file in a separate output folder ----
setwd("set/output/directory")
write.csv(lt,"raw_data_assay2_census.csv")
lt <- read.csv("raw_data_assay2_census.csv")

#setting something as a factor
lt$exp_code = as.factor(lt$exp_code)
lt$nests = as.factor (lt$nests)

## Creating dataset per age for analysis ----
wol1 <- data.frame(subset (lt, time == "one month"))
wol2 <- data.frame(subset (lt, time == "two months"))
wol3 <- data.frame(subset (lt, time == "three months"))
wol4 <- data.frame(subset (lt, time == "four months"))
wol5 <- data.frame(subset (lt, time == "five months"))
wol6 <- data.frame(subset (lt, time == "six months"))
wol7 <- data.frame(subset (lt, time == "seven months"))

# creating data set from 1-4 month and 4-7 month
wol14 <- rbind(wol1,wol2,wol3, wol4)# late-instar larvae have been counted
wol57 <- rbind(wol5,wol6,wol7) # late-instar larvae where not counted
wol47 <- rbind(wol4,wol5,wol6,wol7)# colonies are reproducing


# ------------------ Functions for evaluating GLM and producing test statistics of ALL models -------
# Goodness of fit for GLM ----
# Chi-square test on the residual deviance and degree of freedon
# ran GLM with Poisson, quasipoisson and negative binomial model
# null hypothesis = family (Poisson, quasipoisson or negative binomial) is an adquate fit for the data
# if P > 0.05, family specified in the model is a good fit, i.e., accept null hypothesis
p.gof <- function(model){
  dev <- deviance(model)
  rdf <- df.residual(model)
  pchisq(dev,
         df=rdf,
         lower.tail=FALSE)
}

# Overdispersion of GLM -----
overdisp_fun <- function(model) {
  rdf <- df.residual(model)
  rp <- residuals(model,type="pearson")
  Pearson.chisq <- sum(rp^2)
  prat <- Pearson.chisq/rdf
  pval <- pchisq(Pearson.chisq, df=rdf, lower.tail=FALSE)
  c(chisq=Pearson.chisq,ratio=prat,rdf=rdf,p=pval)
}

# if overdispersed then use quasipoisson or negative binomial
# selected between the 2 by looking at 3 criteria in order of preference -
#(a) linear fit of residuals (q-q plot), (b) goodness of fit of modelm & (c) lack of over dispersion


# Test statistics for model ----
stats.glmer <- function(model){
  drop1(model,.~.,test="Chisq")
}
stats.glm <- function(model){
  drop1(model,.~.,test="F")
}
stats.hdl <- function(model){
  drop1(model,.~.,test="Chisq")
}


# Q-Q plot of model ----
qq <- function(model){
    res <- residuals(model)
    qqnorm(res)
    qqline(res, probs=c(0.25,0.75))
}


#  ------------------ EFFECT OF WOLBACHIA & TIME ON COLONY LIFE HISTORY TRAITS ----------------
# GLMM 1:testing fixed effect with nested random effect -> to test overall effect of predictor variables in a repeated measure model
# GLM 2:testing effect of interaction between predictor variables (no random factor)
# GLM : Age specific analysis. NO random factor, since sr_colony (queen source colony) is not repeatedly measured within an age.
# Chi-square test of each model to produce test statistics presented in paper and graphs.

## Late-instar larvae in colonies ----
#using data collected from 1 to 4 months after starting the assay
ol.fixed <- glmer (old_larvae~wolbachia + time + obs
              + (1|exp_code),
              data = wol14,family=poisson(link=log),
              control=glmerControl(optimizer="bobyqa",
                                   optCtrl=list(maxfun=3e5)))
model <- ol.fixed
summary(model) 
stats.glmer(model) 
qq(model)

ol.intr.nb <- glm.nb (old_larvae~wolbachia + time 
                + wolbachia:time + obs , 
                data = wol14)
model <- ol.intr.nb
p.gof(model) #p = 0.14
overdisp_fun(model) #p = 0.74
summary(model) 
stats.glm(model) 
qq(model)

# age-specific models
#one month
ol.1m.nb <- glm.nb (old_larvae~wolbachia  + obs + queens, 
              data = wol1)
model <- ol.1m.nb
p.gof(model) #p = 0.22
overdisp_fun(model) #p = 0.23
summary(model) 
stats.glm(model) 
qq(model)

#two months
ol.2m.nb <- glm.nb (old_larvae~wolbachia  + obs + queens, 
                data = wol2)
model <- ol.2m.nb
p.gof(model) #p =0.167
overdisp_fun(model)#p =0.47
summary(model) 
stats.glm(model) 
qq(model)

#three months
ol.3m.nb <- glm.nb (old_larvae~wolbachia + obs + queens, 
                 data = wol3)
model <- ol.3m.nb
p.gof(model) #p = 0.19
overdisp_fun(model) #p = 0.37
summary(model) 
stats.glm(model) 
qq(model)

#four months
ol.4m.nb <- glm.nb (old_larvae~wolbachia + queens, 
                 data = wol4)
model <- ol.4m.nb
p.gof(model) #p = 0.26
overdisp_fun(model) #p = 0.44
summary(model) 
stats.glm(model) 
qq(model)

## Eggs in colonies  ----
eg.fixed <- glmer (eggs~wolbachia + time + log(old_larvae) 
                + obs + (1|exp_code),
                data = wol14,family=poisson(link=log),
                control=glmerControl(optimizer="bobyqa",optCtrl=list(maxfun=3e5)))
model <- eg.fixed
summary(model) 
stats.glmer(model) 
qq(model)

eg.intr.nb <- glm.nb (eggs~wolbachia + time + wolbachia:time 
                +log(old_larvae) + obs + queens, 
                data = wol14)
model <- eg.intr.nb
p.gof(model) #p = 0.14
overdisp_fun(model) # p= 0.18
summary(model) 
stats.glm(model)
qq(model)

#age-specific models
# used Poisson, quasi-poisson and negative binomial models to test which one is a good fit

# One month - selected negative binomial
eg.1.nb <- glm.nb (eggs~wolbachia + log(old_larvae)+
                  obs+ queens, 
                data = wol1)
model <- eg.1.nb
p.gof(model) #p =0.2
overdisp_fun(model) #p =0.19
summary(model) 
stats.glm(model) 
qq(model)

# two months - selected negative binomial
eg.2.nb <- glm.nb (eggs~wolbachia + log(old_larvae)+
                     obs+ queens, 
                   data = wol2)
model <- eg.2.nb
p.gof(model) #p =0.16
overdisp_fun(model) #p = 0.16
summary(model) 
stats.glm(model) 
qq(model)

# three months - selected negative binomial
eg.3.nb<- glm.nb (eggs~wolbachia + log(old_larvae)+
                  obs+ queens, 
                data = wol3)
model <- eg.3.nb
p.gof(model) #p = 0.11
overdisp_fun(model) #p = 0.15
summary(model) 
stats.glm(model) 
qq(model)

# four months - selected negative binomial
eg.4.nb<- glm.nb (eggs~wolbachia + log(old_larvae)+ queens, 
                  data = wol4)
model <- eg.4.nb
p.gof(model) #p =0.23
overdisp_fun(model) # p = 0.30
summary(model) 
stats.glm(model) 
qq(model)

## Worker pupae in colonies ----
# when late-instar larvae were counted
# data analysed exactly like net productivity
wk.fixed.14 <- glmer (worker_pup~wolbachia + time 
                      +log(old_larvae) + log(queens) + obs
                      + (1|exp_code),
                      data = wol14,family=poisson(link=log),
                      control=glmerControl(optimizer="bobyqa",
                                           optCtrl=list(maxfun=3e5)))
model <- wk.fixed.14
summary(model) 
stats.glmer(model) 
qq(model)

# worker_pupuctivity when late-instar larvae were not counted
wk.fixed.57 <- glmer (worker_pup~wolbachia + time + log(queens)
                      + (1|exp_code),
                      data = wol57,family=poisson(link=log),
                      control=glmerControl(optimizer="bobyqa",
                                           optCtrl=list(maxfun=3e5)))
model <- wk.fixed.57
summary(model) 
stats.glmer(model) 
qq(model)

# effect of interaction term when late-instar larvae were counted
wk.intr.14.qp <- glm (worker_pup~wolbachia + time + 
                        wolbachia:time 
                      + log(old_larvae) + obs + queens, 
                      data = wol14,family=quasipoisson(link=log))
model <- wk.intr.14.qp
summary(model) 
stats.glm(model) 
qq(model)

# effect of interaction term when late-instar larvae were not counted
wk.intr.57.nb <- glm.nb (worker_pup~wolbachia + time +
                           wolbachia:time + queens, 
                         data = wol57)
model <- wk.intr.57.nb
p.gof(model) #p = 0.18
overdisp_fun(model) #p = 0.42
summary(model) 
stats.glm(model) 
qq(model)

# time specific
# one month
wk.1.nb <- glm.nb (worker_pup~wolbachia + log(old_larvae)
                   + obs+ queens, 
                   data = wol1)
model <- wk.1.nb
p.gof(model) #p = 0.14
overdisp_fun(model) #p = 0.39
summary(model) 
stats.glm(model) 
qq(model)

# two months
wk.2.nb <- glm.nb (worker_pup~wolbachia + log(old_larvae)
                   + obs+ queens, 
                   data = wol2)
model <- wk.2.nb
p.gof(model) #p = 0.15
overdisp_fun(model) #p =0.165
summary(model) 
stats.glm(model) 
qq(model)

# three months
wk.3.nb <- glm.nb (worker_pup~wolbachia + log(old_larvae)
                   + obs+ queens, 
                   data = wol3)
model <- wk.3.nb
p.gof(model) #p = 0.11
overdisp_fun(model) #p =0.13
summary(model) 
stats.glm(model) 
qq(model)

# four months
wk.4.nb <- glm.nb (worker_pup~wolbachia + log(old_larvae)
                   + queens, 
                   data = wol4)
model <- wk.4.nb
p.gof(model)#p =0.23
overdisp_fun(model) #p = 0.31
summary(model) 
stats.glm(model) 
qq(model)

# Five months
wk.5.nb <- glm.nb (worker_pup~wolbachia + queens, 
                   data = wol5)
model <- wk.5.nb
p.gof(model) #p =0.26
overdisp_fun(model) #p = 0.19
summary(model) 
stats.glm(model) 
qq(model)

# Six months
wk.6.nb <- glm.nb (worker_pup~wolbachia + queens, 
                   data = wol6)
model <- wk.6.nb
p.gof(model) #p =0.25
overdisp_fun(model) #p = 0.42
summary(model) 
stats.glm(model) 
qq(model)

# Seven months
wk.7.nb <- glm.nb (worker_pup~wolbachia + queens, 
                   data = wol7)
model <- wk.7.nb
p.gof(model) #p =0.24
overdisp_fun(model) #p = 0.42
summary(model) 
stats.glm(model) 
qq(model)

## Queens in colonies ----
# effect when some colonies start reproducing
qn.fixed47 <- glmer (tot_qn~wolbachia + time + (1|exp_code),
                   data = wol47,family=poisson(link=log),
                   control=glmerControl(optimizer="bobyqa",optCtrl=list(maxfun=3e5)))
model <- qn.fixed47
summary(model) 
stats.glmer(model) 
qq(model)

qn.intr.nb <- glm.nb (tot_qn~wolbachia + time + wolbachia:time , 
                      data = wol47)
model <- qn.intr.nb
p.gof(model) #p =0.1
overdisp_fun(model) #p = 0.21
summary(model) 
stats.glm(model) 
qq(model)

#Age specific effects
# four months
qn.4.nb <- glm.nb (tot_qn~wolbachia + log(old_larvae), 
                   data = wol4)
model <- qn.4.nb
p.gof(model) #p =0.28
overdisp_fun(model) #p = 0.3
summary(model) 
stats.glm(model) 
qq(model)

# Five months
qn.5.nb <- glm.nb (tot_qn~wolbachia , 
                   data = wol5)
model <- qn.5.nb
p.gof(model) #p =0.24
overdisp_fun(model) #p = 0.1
summary(model) 
stats.glm(model) 
qq(model)

# Six months
qn.6.nb <- glm.nb (tot_qn~wolbachia , 
                   data = wol6)
model <- qn.6.nb
p.gof(model) #p =0.22
overdisp_fun(model) #p = 0.71
summary(model) 
stats.glm(model) 
qq(model)

# Seven Months
qn.7.nb <- glm.nb (tot_qn~wolbachia , 
                   data = wol7)
model <- qn.7.nb
p.gof(model) #p =0.21
overdisp_fun(model) #p = 0.52
summary(model) 
stats.glm(model) 
qq(model)

## Males in colonies ----
# working with data where colonies started reproducing
ml.fixed <- glmer(tot_male~wolbachia + time+
                    (1|exp_code),
                  data = wol47,family = poisson(link = log))
model <- ml.fixed
summary(model) 
stats.glmer(model) 
qq(model)
stats.glmer(model)

ml.intr.nb <- glm.nb (tot_male~wolbachia + time
                      + wolbachia:time,data = wol47)
model <- ml.intr.nb
p.gof(model) #p = 0.97
overdisp_fun(model) #p = 0.99
summary(model) 
stats.glm(model) 
qq(model)

# Time specific effects of Wolbachia
# Four months
ml.4.hdl <- hurdle (tot_male~wolbachia, 
                  data = wol4, dist = "negbin")
model <- ml.4.hdl
summary(model)
stats.hdl(model)

# Five months
ml.5.hdl <- hurdle (tot_male~wolbachia, 
                    data = wol5, dist = "negbin")
model <- ml.5.hdl
summary(model)
stats.hdl(model)

# Six months
ml.6.hdl <- hurdle (tot_male~wolbachia, 
                    data = wol6, dist = "negbin")
model <- ml.6.hdl
summary(model)
stats.hdl(model)

# Seven months
ml.7.hdl <- hurdle (tot_male~wolbachia, 
                    data = wol7, dist = "negbin")
model <- ml.7.hdl
summary(model)
stats.hdl(model)


# ------------------- Dry Mass differences across time and Wolbachia infection -----
# processing raw data ----
dm$exp_code <- as.factor(dm$exp_code)
dm$sample <- as.factor(dm$sample)

#check disrtibution of data
hist(dm$mass) #normally distributed, thus use LM and LMM

## dry mass: Creating age specific datasets ----
dm.2 <- data.frame(subset (dm, time == "two months"))
dm.3 <- data.frame(subset (dm, time == "three months"))
dm.4 <- data.frame(subset (dm, time == "four months"))
dm.6 <- data.frame(subset (dm, time == "six months"))

# Linear Mixed Effect Models for dry mass -----
#repeated measures since colonies analysed over a period of time.
# Model : LMM to assess FIXED and INTERACTION effects, with random factors.
# Model 'time specific': LMM for each time point
# used ANOVA to produce test statistic
dm.lmm <- lmer (log(mass)~wolbachia*time + (1|exp_code),
                 data = dm)
model = dm.lmm
car::Anova(model, ddf="Satterthwaite")
res <- residuals(model)
shapiro.test(res)
qq(model)

#producing statistics for multiple pairwise comparisons of dry mass over time
summary(glht(dm.fixed, linfct= mcp(time = "Tukey"))) 

# experimental colonies have been used as random factors as there are multiple measurements per colony (samples)
# Worker dry mass after 2 months of starting the assay
lm.dm.2 <- lmer (log(mass)~wolbachia+(1|exp_code),
                  data = dm.2)
car::Anova(lm.dm.2, ddf="Satterthwaite")
res <- residuals(lm.dm.2)
shapiro.test(res) #normally distributed

# Worker dry mass after 3 months of starting the assay
lm.dm.3<- lmer (log(mass)~wolbachia + (1|exp_code),
                  data = dm.3)
car::Anova(lm.dm.3, ddf="Satterthwaite")
res <- residuals(lm.dm.3) #normally distributed
shapiro.test(res)

# Worker dry mass after 4 months of starting the assay
lm.dm.4 <- lmer (log(mass)~wolbachia + (1|exp_code),
                  data = dm.4)
car::Anova(lm.dm.4, ddf="Satterthwaite")
res <- residuals(lm.dm.4)
shapiro.test(res) #normally distributed

# Worker dry mass after 6 months of starting the assay
lm.dm.6 <- lmer (log(mass)~wolbachia + (1|exp_code),
              data = dm.6)
car::Anova(lm.dm.6, ddf="Satterthwaite")
res <- residuals(lm.dm.6)
shapiro.test(res)


# --------------------- Plotting graphs -----------------------
# Fig 3 : mean and 95% CI of trait value ----
# each graph produced independently and then compiled in Inkscape
# use this to show change over time
my.color <-c ("chocolate2",  "darkorchid4")

lt$m <- lt$tot_male #creating a universal variable for plotting
dm$m <- dm$mass

#summarizing the statistics for trait
mean <- summarySE(lt, measurevar="m", 
                  groupvars=c("wolbachia", "time"), na.rm = T)

#re-ordering variables as desired
mean$time <- factor(mean$time, levels= c("one month","two months" ,"three months",
                                         "four months","five months", "six months",
                                         "seven months"))
mean$wolbachia<- factor(mean$wolbachia, 
                        levels = c("uninfected", "infected"))

# generating a custom plot. Texts were later edited in Inkscape
ggplot(mean, aes(x=time,y = m, group = wolbachia)) + 
  scale_colour_manual(values = my.color)+
  scale_fill_manual(values = my.color)+
  geom_errorbar(aes(ymin= m-ci, ymax= m+ci, colour = wolbachia),
                linetype = "longdash", 
                size = 0.5, width= 0.1, 
                alpha = 0.8,position= position_dodge(0.2))+
  geom_line(aes(color = wolbachia),
            size = 0.8,position= position_dodge(0.2))+
  geom_point(aes(fill = wolbachia),shape = 21, 
             size = 4, position = position_dodge(0.2))+
  ggtitle("Worker Pupae Mass")+
  theme(strip.background = element_blank(), strip.placement = "outside",
        strip.text = element_text(family = "Arial", colour = "black", face = "italic",size = "12"))+
  theme(plot.title = element_text(hjust = 0.5))+ 
  labs(y = "adult and pupae count") +
  theme(panel.background=element_rect(fill = NA),
        plot.background=element_rect(fill = NA),
        panel.border = element_rect(colour = "black", fill = NA, size = 0.8),
        panel.grid.major = element_blank(),
        panel.grid.minor = element_blank(),
        legend.position = "none",
        #legend.position = c(0.08,0.80),
        legend.text = element_text(family = "Arial", colour = "black", size = "10"),
        legend.background = element_rect(fill = NA),
        legend.key = element_rect(color = NA, fill = NA),
        axis.title = element_text(family = "Arial", colour ="black", face = "bold", size = "11"),
        axis.text = element_text(family = "Arial", colour = "black", size = "12"),
        axis.title.x = element_blank(),
        axis.ticks = element_line(colour = "black", size = 1),
        plot.title = element_text(family = "Arial", colour = "black", face = "bold", size = "14", 
                                  margin = margin(t='1', r='0.5', b='0.5', l='1', unit = 'cm')))
#saving image in desired format to the file path set in the start of the script. Default saves last plot.
ggsave(file = "2w_mass.png", dpi = 300)

# Supplementary : Box plots for dry mass -----
my.color <-c ("chocolate2",  "darkorchid4")
dm$m <- dm$mass
dm$time <- factor(dm$time, levels= c("two months", "three months", 
                                     "four months","six months"))
dm$wolbachia<- factor(dm$wolbachia, 
                       levels = c("uninfected", "infected"))

ggplot(dm, aes(x=time,y = m, fill = wolbachia)) + 
  geom_boxplot(notch = T,position= position_dodge(0.8))+
  #scale_colour_manual(values = my.color)+
  scale_fill_manual(values = my.color)+
  ggtitle("Worker Pupae Mass")+
  theme(strip.background = element_blank(), strip.placement = "outside",
        strip.text = element_text(family = "Arial", colour = "black", face = "italic",size = "12"))+
  theme(plot.title = element_text(hjust = 0.5))+ #adjusts this to middle
  labs(y = "dry  mass (mg.)") +
  theme(panel.background=element_rect(fill = NA),
        plot.background=element_rect(fill = NA),
        panel.border = element_rect(colour = "black", fill = NA, size = 0.8),
        panel.grid.major = element_blank(),
        panel.grid.minor = element_blank(),
        legend.position = "none",
        #legend.position = c(0.08,0.80),
        legend.text = element_text(family = "Arial", colour = "black", size = "10"),
        legend.background = element_rect(fill = NA),
        legend.key = element_rect(color = NA, fill = NA),
        axis.title = element_text(family = "Arial", colour ="black", face = "bold", size = "11"),
        axis.text = element_text(family = "Arial", colour = "black", size = "12"),
        axis.title.x = element_blank(),
        axis.ticks = element_line(colour = "black", size = 1),
        plot.title = element_text(family = "Arial", colour = "black", face = "bold", size = "14", 
                                  margin = margin(t='1', r='0.5', b='0.5', l='1', unit = 'cm')))
#saving image in desired format to the file path set in the start of the script. Default saves last plot.
ggsave(file = "2w_mass_box.png", dpi = 300)
